# The early life microbiota protects neonatal mice from pathological small intestinal epithelial cell shedding

**DOI:** 10.1101/789362

**Authors:** Kevin R Hughes, Z Schofield, MJ Dalby, S Caim, L Chalklen, F Bernuzzi, C Alcon-Giner, G Le Gall, AJM Watson, LJ Hall

## Abstract

The gut microbiota plays a crucial role in regulating and maintaining the epithelial barrier, particularly during early life. Notably, patients with chronic intestinal inflammation have a dysregulated process of renewal and replenishment of the intestinal epithelial cell (IEC) barrier, which is linked to disturbances in the gut microbiota. To date, there are no studies focussed on understanding the impact of inflammatory cell shedding events during the early life developmental window, and which host and microbial factors mediate these responses. Here we sought to determine pathological cell shedding outcomes throughout the postnatal developmental period (day 14, 21, 29 and week 8). Surprisingly neonatal mice (day 14 and 21) were highly refractory to induction of cell shedding after intraperitoneal administration of LPS, with day 29 mice showing strong pathological responses, more similar to those observed in adult mice. These differential responses were not linked to defects in the cellular mechanisms and pathways known to regulate cell shedding responses, although we did observe that neonatal mice had elevated anti-inflammatory (IL-10) responses. Notably, when we profiled microbiota and metabolites from these mice, we observed significant alterations. Neonatal mice had high relative abundances of *Streptococcus*, *Escherichia* and *Enterococcus* and increased primary bile acids. In contrast, older mice were dominated by *Candidatus* Arthromitus, *Alistipes* and *Lachnoclostridium*, and had increased concentrations of SCFAs and methyamines. Faecal microbiota transplant (FMT) and antibiotic studies confirmed the importance of early life gut microbiota in cell shedding responses. In these studies, neonates treated with antibiotics restored LPS-induced small intestinal cell shedding, whereas adult FMT alone had no effect. Our findings further support the importance of the early life window for microbiota-epithelial interactions in the presence of inflammatory stimuli and highlight areas for further investigation to probe underlying mechanisms to drive therapeutic development within the context of chronic inflammatory intestinal diseases.

## Introduction

Early life represents a critical developmental window, which lays the foundations for short- and long-term health [1, 2]. Immediately after birth, the human gastrointestinal (GI) tract is colonised by a succession of microbes, up until 2-3 years of age when the ‘adult’ microbiota fully establishes. This early life ecosystem is relatively simple in composition, with *Bifidobacterium* the dominant bacterial genus, which is linked to their ability to digest breast-milk derived dietary components (i.e. human milk oligosaccharides), with *Bacteroides*, *Veillonella*, *Faecalibacterium*, *Streptococcus*, and *Escherichia* also important microbiota members [1, 3]. Notably, several factors influence the development of the early life microbiome, including mode of delivery, host genetics, diet and drugs (e.g. antibiotics), with disturbances linked to an increasing number of immune-linked diseases including ulcerative colitis (UC), arthritis, and atopic allergy [2, 4].

During early life the gut microbiota plays a key role in development and maturation of numerous host responses [5]. A central interface mediating these interactions is the intestinal lumen, which represents a unique ecological environment that is the major site for microbial-host crosstalk. Indeed, recent studies indicate that the microbiota, and certain microbiota members, modulate transcriptional and functional intestinal epithelial cell (IEC) responses through pathological and immune-mediated mechanisms [6, 7]. A central output of these interactions is the reinforcement of the gut barrier, which is achieved through constant replenishment of cells [8]. Intestinal epithelial cells are ‘born’ in the crypts of Lieberkuhn from where they migrate to the tips of the intestinal villi and are lost into the intestinal lumen through a homeostatic process known as cell shedding [9, 10]. Cell shedding represents an important host physiological response which ensures continuity of the intestinal epithelial layer despite continuous replacement of intestinal epithelial cells every 3-5 days [11]. Crucially, intestinal inflammation is correlated with elevated or ‘pathological’ cell shedding, which is also correlated with altered microbiota profiles [12–14]. Previous studies have indicated that antibiotic usage, and antibiotic-induced microbiota perturbations, particularly in infancy, are linked to increased risk of developing inflammatory disorders in later life [15]. More recently, research, including our own, has shown important links between members of the intestinal microbiota and the cell shedding response *in vivo*, demonstrating that early life members of the gut microbial community (i.e. *Bifidobacterium*) can drive protection against pathological cell shedding in adult mice [16, 17].

Although previous studies have demonstrated that there are numerous microbiota-mediated effects on IECs during neonatal development [18], there are no studies focussed on the role of early life gut microbiota during pathological cell shedding. Here, we sought to determine how postnatal development stage impacts pathological cell shedding using an established *in vivo* model and using histological, immune, and microbial/metabolite profiling to assess intestinal barrier function in early life and understand the mechanisms involved in regulating this response. Intriguingly, we observed a protective effect in neonatal mice during LPS-induced epithelial shedding when compared to adult counterparts, which was diminished after faecal microbiota transplant (FMT) and antibiotic-induced microbiota disturbances, highlighting the crucial role the microbiota plays during this important early life period.

## Results

### Neonatal mice are refractory to LPS-induced pathological cell shedding

We previously demonstrated that IP administration of LPS induced potent cell shedding in adult mice with peak response 90 minutes post injection as determined by cleaved caspase-3 (CC3) staining [16, 19]. To assess cell shedding response of neonatal mice, 14 and 21 day old mice were administered with either low (1.25mg kg-1) or high (10mg kg-1) dose LPS (or PBS vehicle control) and sacrificed 90 minutes post-administration (Figure 1A). Surprisingly, intestinal epithelial cells (IECs) in day 14 and 21 old mice were CC3 negative following low and high dose LPS and villi exhibited normal histology at the macroscopic level (Figure 1B and C). We next treated 29-day old neonatal mice with IP LPS and observed the re-emergence of cell shedding response (Figure 1B and D). As expected, adult mice (8-10 weeks) showed enhanced cell shedding responses as determined by CC3 positive cells and associated pathology within the small intestine (Figure 1B and E).

**Figure 1:**
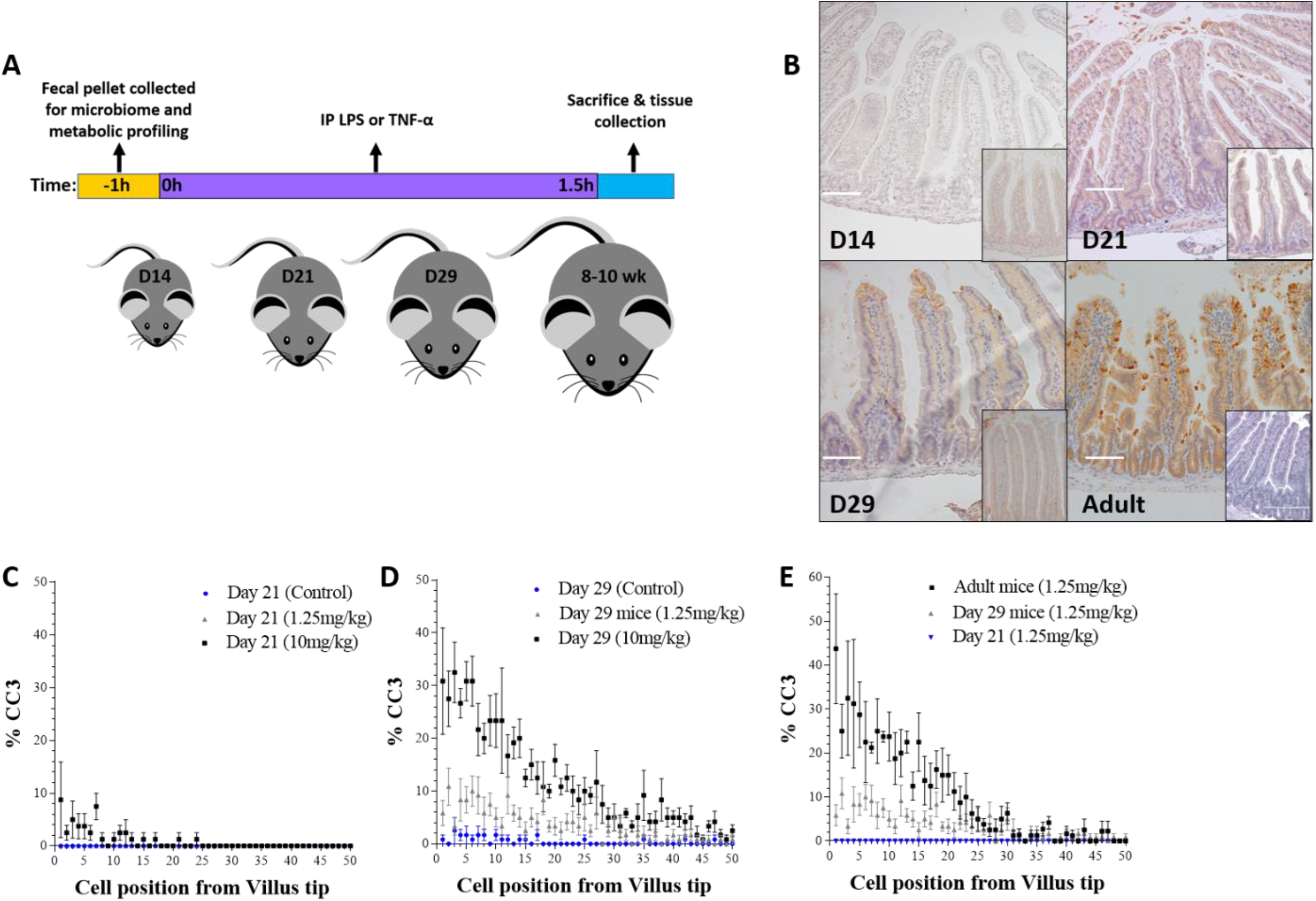
Neonatal mice are refractory to LPS-induced cell shedding. (**A**) Experimental workflow for LPS/TNF-alpha administration and tissue collection. (**B**) Immuno-histochemical analysis of cleaved-caspase 3 (CC3) on haematoxylin stained intestinal tissue demonstrated absence of cell shedding at day 14 with only occasional cell shedding events observed at day 21. Mice at day 29 demonstrated emergence of cell shedding in response to LPS administration. Inset images show representative control tissue. (**C-E**) Wincrypts software was used to quantitate cell shedding events in 50 representative half villi, shown as percentage CC3 positive versus cell position from the villus tip. Significant difference in rate of observed cell shedding events (P<0.0001) was found between adult and neonatal tissue. White scale bar 100 μm.

We reasoned that absence of CC3 staining in neonates may be the result of delayed IEC apoptosis. To test this hypothesis, 14-day old neonatal mice were administered low or high dose LPS and tissue collected up to 180 minutes post-LPS administration. In contrast to adult mice which show extensive CC3 staining at 90 minutes coupled with associated villus atrophy at later time points, neonatal mice showed no evidence of CC3 staining up to 180 minutes post-LPS, nor evidence of pathological changes that normally accompany the cell shedding response (data not shown). Taken together, mice are largely refractory to LPS induced cell shedding during the neonatal period, with intermediate responses observed in day 29 old mice, with adult mice very susceptible to pathological cell shedding. These distinctions cannot be explained by altered kinetics of cell death responses during the early life developmental period.

### Cell signalling pathways that drive pathological cell shedding are expressed in neonatal mice

Previous work suggests that systemically delivered LPS signals via mononuclear cells which express TLR4. This signalling cascade is mediated by the TLR-adaptor protein MYD88, which drives production of TNF-ɑ, which in turn binds TNFR1 expressed on the surface of IECs and triggers apoptosis and cell shedding [19]. We reasoned that this signalling cascade may be diminished in neonatal mice, thus preventing initiation of the cell shedding response. IECs were isolated from the mucosa of neonatal (day 14), and adult mice and expression of TNF-R1 determined by real-time PCR. Levels of TNFR1 transcript was significantly increased in control neonatal mice compared to adult mice, but neonates dosed with LPS had significantly reduced TNFR1 (Figure 2A). We next reasoned that production of TNF-ɑ in neonatal mice may be impaired. However, ELISA analysis of whole small intestinal homogenates from day 14 neonatal mice exposed to low and high dose LPS confirmed production of TNF-ɑ at levels comparable to adult mice (Figure 2B).

**Figure 2:**
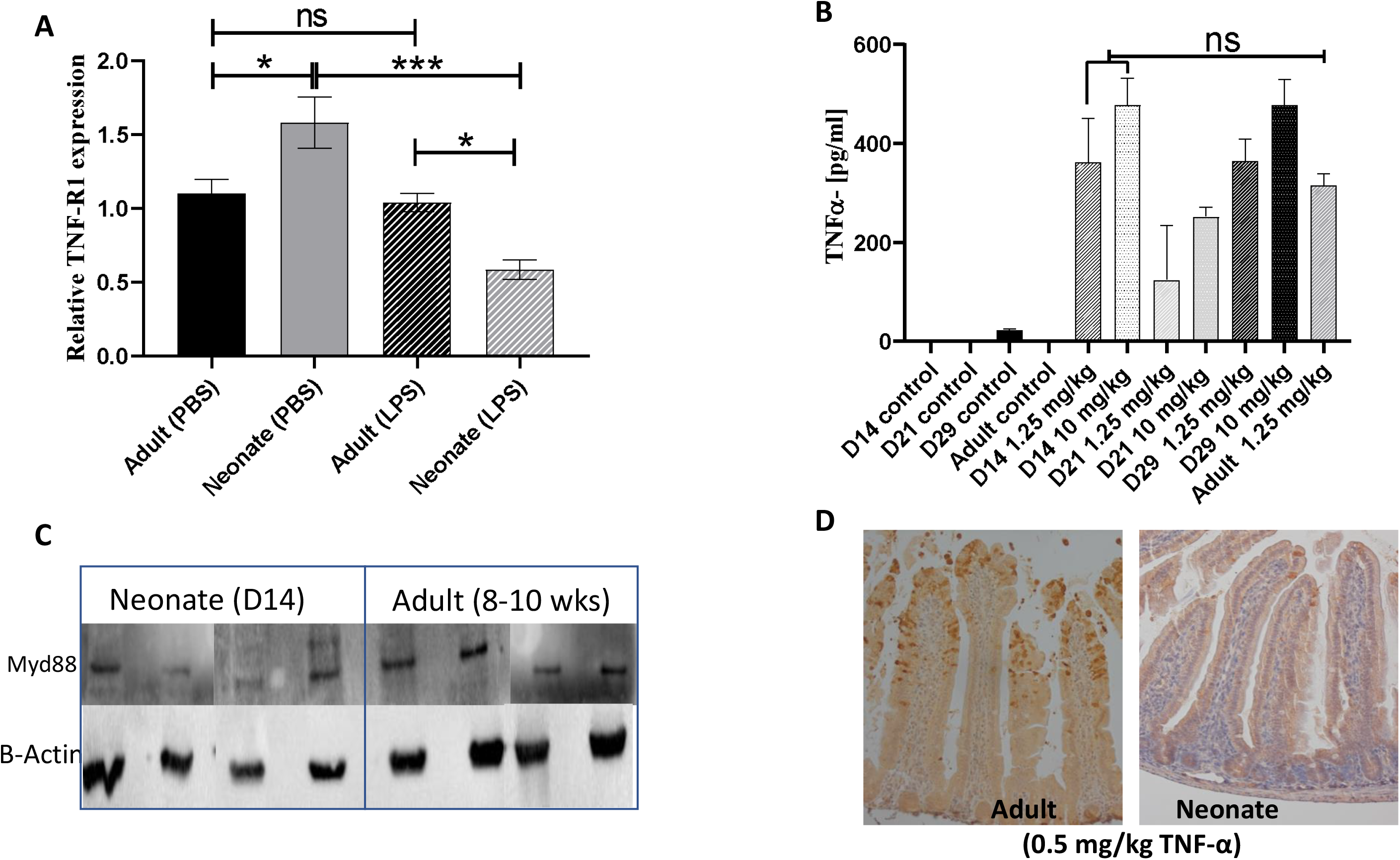
Neonates do not have impaired TNF signalling pathways. Expression and protein quantification was used to investigate the molecular basis of differential cell shedding responses of neonatal and adult mice. (**A**) TNF-R1 expression was quantified by real-time RT-PCR in isolated intestinal epithelial cells and showed control neonates had higher expression (with the opposite observed in LPS-treated mice). (**B**) ELISA analysis of TNF-α expression in whole small intestinal homogenates showed no significant difference between induction of TNF-α in day 14 versus adult mice. (**C**) Western blot analysis of MYD88 protein in murine spleen tissue confirmed similar levels of expression in neonatal (day 14) and adult mice. (**D**) Challenge with IP TNF-alpha (0.5mg / kg) confirmed that neonatal mice were largely refractory to TNF-α induced cell shedding versus adult counterparts. Statistical analysis performed by one way anova with Tukeys post test (n = 5±2 ***p<0.001, *p<0.05, ns = not significant).

Previous studies have shown reduced expression of MYD88 protein in human neonatal peripheral blood leukocytes, and we hypothesised that altered MYD88 expression in neonatal mice may drive reduced cell shedding [20]. However, there were no significant differences in the levels of MYD88 protein in homogenised spleen samples taken from 14 day old and adult mice (Figure 2C). To confirm our findings indicating there are no neonatal defects in cell shedding signalling pathways we also challenged mice IP with TNF-ɑ. Whilst adult mice showed the expected potent cell shedding response, neonatal mice were largely refractory to cell shedding, with only isolated cell shedding events observed (Figure 2D). Taken together, this suggests that core components of the cell shedding signalling pathways are intact in neonatal mice and that blunting of the cell shedding response cannot be explained by altered expression profiles of these components at the transcriptional and translational level.

### Inflammatory responses are altered in neonatal versus adult mice during pathological cell shedding

PCR and antibody arrays were used to explore if differing cell shedding outcomes in neonatal mice may be driven by alterations in immunologic/inflammatory responses and/or in expression of bacterial recognition markers. We observed reduced expression of pro-inflammatory markers such as IFN-γ but increased anti-inflammatory cytokines such as IL-10 in neonatal mice versus adults following LPS stimulation (Figure 3A). We also observed altered expression of transcripts associated with bacterial recognition including nucleotide-binding oligomerization domain-like receptors (NODs) and members of TNF-receptor associated proteins including TRAF1, TNFrsf10 and TNFrsf21, alongside changes to components of the apoptotic cascade including reduced expression of CARD9, 10, 11 and Casp3 (Figure 3B). At a translational level we observed elevation of IL-10, IL-7, IL-12 and TNF-ɑ in neonatal versus adult mice during pathological cell shedding, with the increased IL-10 levels in neonatal mice confirmed by ELISA (**Supplementary Figure 1** and Figure 3C). These data indicate that neonatal mice may have a more anti-inflammatory and anti-apoptotic milieu when compared to their adult counterparts, which may be in part driving altered cell shedding responses.

**Figure 3:**
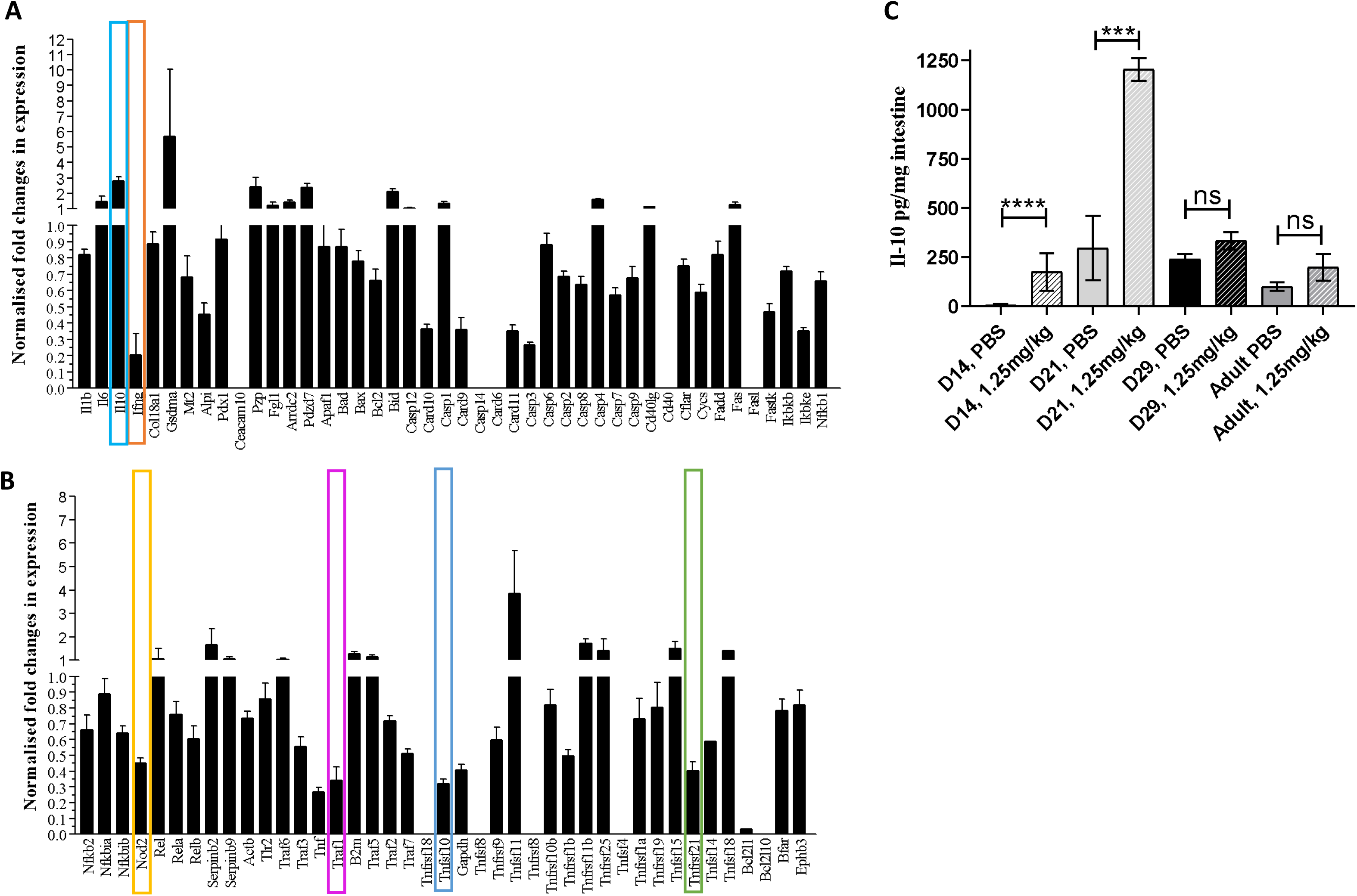
Neonatal mice have altered immune and apoptosis pathways. PCR arrays were used to assess global transcriptional changes in neonatal versus adult intestinal epithelial cells using a custom panel of epithelial / cell shedding related targets. Panels (**A**) and (**B**) show the global transcriptional changes observed. Specific signalling molecules where substantive changes in expression were observed and which are discussed in the associated text are marked. We also assessed IL-10 expression in whole small intestinal homogenates by ELISA. Significant elevation of IL-10 in response to LPS challenge was observed in day 14 (p<0.0001) and day 21 neonatal mice (p<0.001), but not in day 29 or adult mice (**C**). Statistical analysis performed by one way anova with Tukeys post test (n = 5±2 ****p<0.0001, ***p<0.001, ns = not significant).

### Microbiota profiles and microbial-derived metabolites are significantly altered in neonatal mice when compared to adults

Previous studies have indicated that the neonatal microbiota significantly influences immune and epithelial responses [21]. Moreover, humans and mice show distinct alterations in microbiota composition during the early life developmental window, and our previous work highlighted the importance of (early life) gut microbes in regulating the cell shedding response. Thus, we sought to determine how microbial profiles diversify at key postnatal time points which also correspond to the observed differences in cell shedding responses.

Faecal metataxonomic bacterial composition was determined by 16S rRNA gene sequencing. Genus level clustering of samples using non-metric multidimensional scaling (NMDS) indicated separation of the microbiota profiles between day 14 from day 21, day 29 and adult mice (PERMANOVA comparisons P = 0.006), while day 21 mice showed a more divergent microbiota that was separate from day 29 mice (PERMANOVA comparison P = 0.006), which clustered closely with adult (maternal) mouse samples (Figure 4A). The clustering of the microbiota composition in day 14 mice was driven by the genus *Streptococcus* while day 21 mice showed a divergent microbiota composition driven in part by *Lactobacillus* abundance. Day 29 mice clustered with the adult samples with the composition driven by genera including *Roseburia*, *Bilophila* and *Alistipes* (Figure 4A).

**Figure 4:**
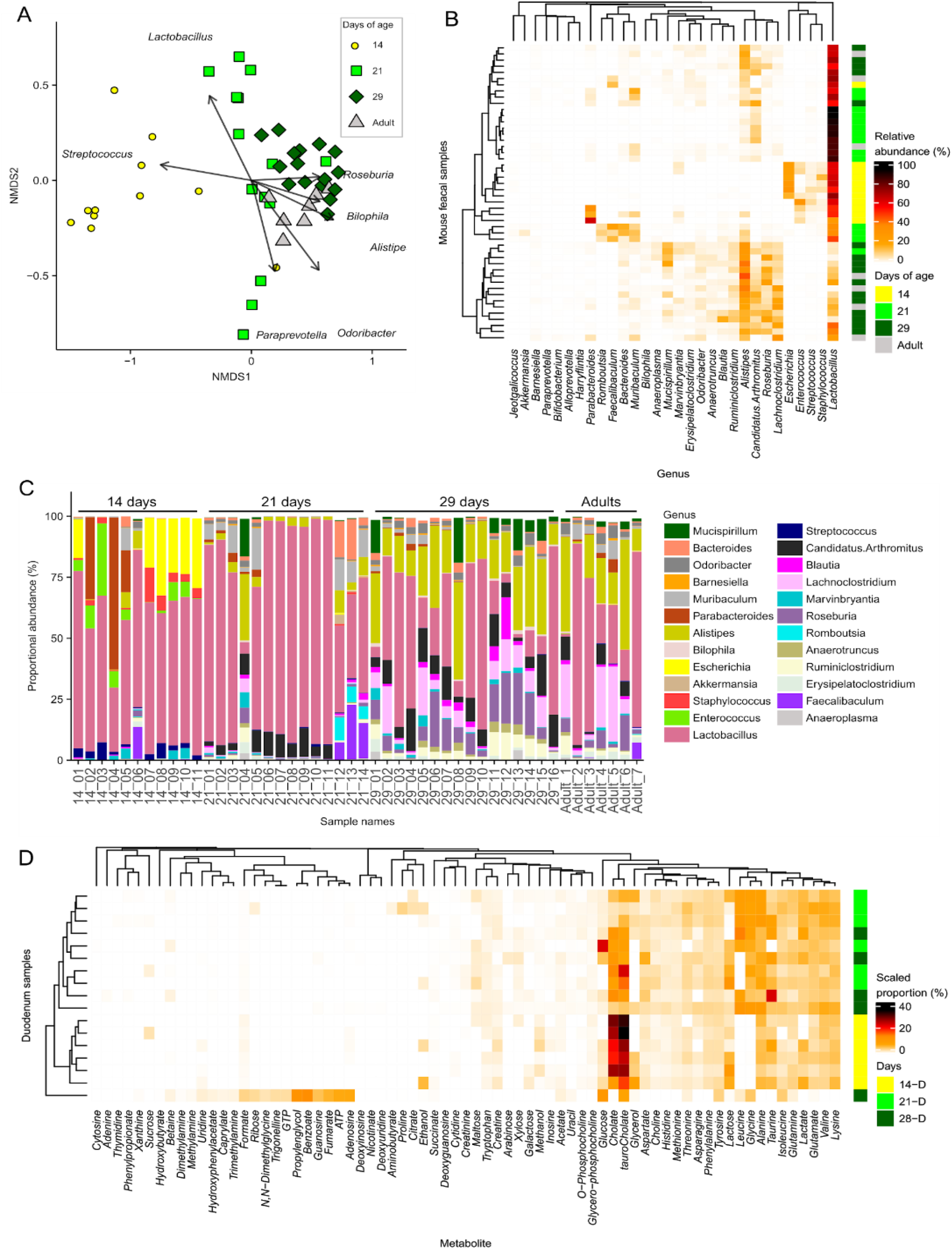
Neonatal mice have altered microbiota genus composition and duodenum metabolite composition. Sequencing analysis of faecal samples taken from mice aged 14, 21, and 29 days and 10 week old adult mice. (**A**) Bray-Curtis-based non-metric multidimensional scaling (NMDS) plot of faecal microbiota samples shows distinct grouping for day 14 and 21 mice but with a distinct shift towards D29 and into adulthood. (**B**) Heatmap shows relative abundance of the 30 most abundant genera clustered using a Bray-Curtis dissimilarity and indicates increased diversity during the weaning period. (**C**) Relative abundance of genera in each sample grouped by day of age. (**D**) Relative abundance of NMR analysis of metabolites in duodenum contents clustered by Bray-Curtis dissimilarity shows distinct grouping for day 14 mice with a greater proportion of cholate and tauro-cholate and an absence of glycine and leucine.

We observed that number of genera detected, and the Shannon diversity index increased during the neonatal period; with the lowest number of genera in day 14 mice compared to day 21, day 29 and adult mice (**Supplementary Fig 2A and 2B**). We also examined the 30 most abundant genera by relative abundance from day 14, 21, 29, and adult mice using Bray-Curtis dissimilarity (Figure 4B), showing that day 14 mice samples clustered with a distinct microbiota composition comprised of *Streptococcus*, *Staphyloccous*, *Escherichia*, and *Enterococcus* that were absent from older samples while *Lachnoclostridium*, *Roseburia*, *Candidatus* Arthromitus (or Segmented filamentous bacteria), and *Alistipes* became dominant by 29 days and in adult mice. This composition was also illustrated with the relative abundance within in mouse sample (Figure 4C). Interestingly, the 21-day old group appeared to have a more varied microbiota profile between animals, possibly linked to the rapidly changing nutritional environment during this period (i.e. moving from maternal milk to solid food) (Figure 4C).

We next performed 1H NMR analysis of duodenal and stomach contents harvested from neonatal and adult mice to profile changes in metabolites (Figure 4D and **Supplementary Figure 2C and D**). Day 14 mice duodenum samples clustered separately with higher proportions of the bile acids cholate and taurocholate (Figure 4D, **Supplementary Figure 2J and 2K)**, at which time the presence of SN-glycerol-phosphocholine in the stomach contents indicated the milk-based diet at 14 days (**Supplementary Figure 2E and 2F)**. Day 14 mice also showed an absence of leucine and glycine from duodenum contents potentially linked to their milk diet (**Supplementary Figure 2I-M)**. The appearance of the microbial metabolites propionate and trimethylamine in stomach contents at day 21 and 28 indicated the start of coprophagy and the potential exposure and transfer of new microbes between neonatal mice. These changes suggested dramatic shifts in microbiota and metabolic profiles during the neonatal period related to changes in diet, which correlates with distinct responses to LPS-induced cell shedding.

### Microbiota composition and antibiotic-induced microbiota disturbances modulate pathological cell shedding responses

Next, we determined if the protective responses observed in neonatal mice were associated with differences in the microbiota and metabolite profiles as seen in Figure 4. We performed an FMT experiment whereby we dosed neonatal mice (day 16) with adult (dam) faecal samples (8-10 weeks) for 5 days via oral gavage, with further groups also treated with antibiotics (via drinking water for 5 days) to deplete the resident neonatal microbiota (alone), and in combination with FMT. On D21 all group received LPS by IP injection (Figure 5A). Histopathological analysis of small intestine showed that adult FMT alone was insufficient to significantly disrupt intestinal pathology (Figure 5B). However, disruption of the neonatal microbiota with antibiotics significantly increased damage to the epithelium, which was not reversed by FMT from adult mice (Figure 5B). Neonatal mice treated with antibiotics (day 10-14) with and without adult FMT had significantly increased apoptotic cell death (measured by CC3 intensity) (Figure 5B and 5C). Although we did not observe a significant increase in intestinal pathology/damage scores from FMT alone, we did observe that FMT induced a significant increase in apoptotic cell death (Figure 5D). These data indicate that the neonatal microbiota plays a key role in modulating cell shedding.

**Figure 5:**
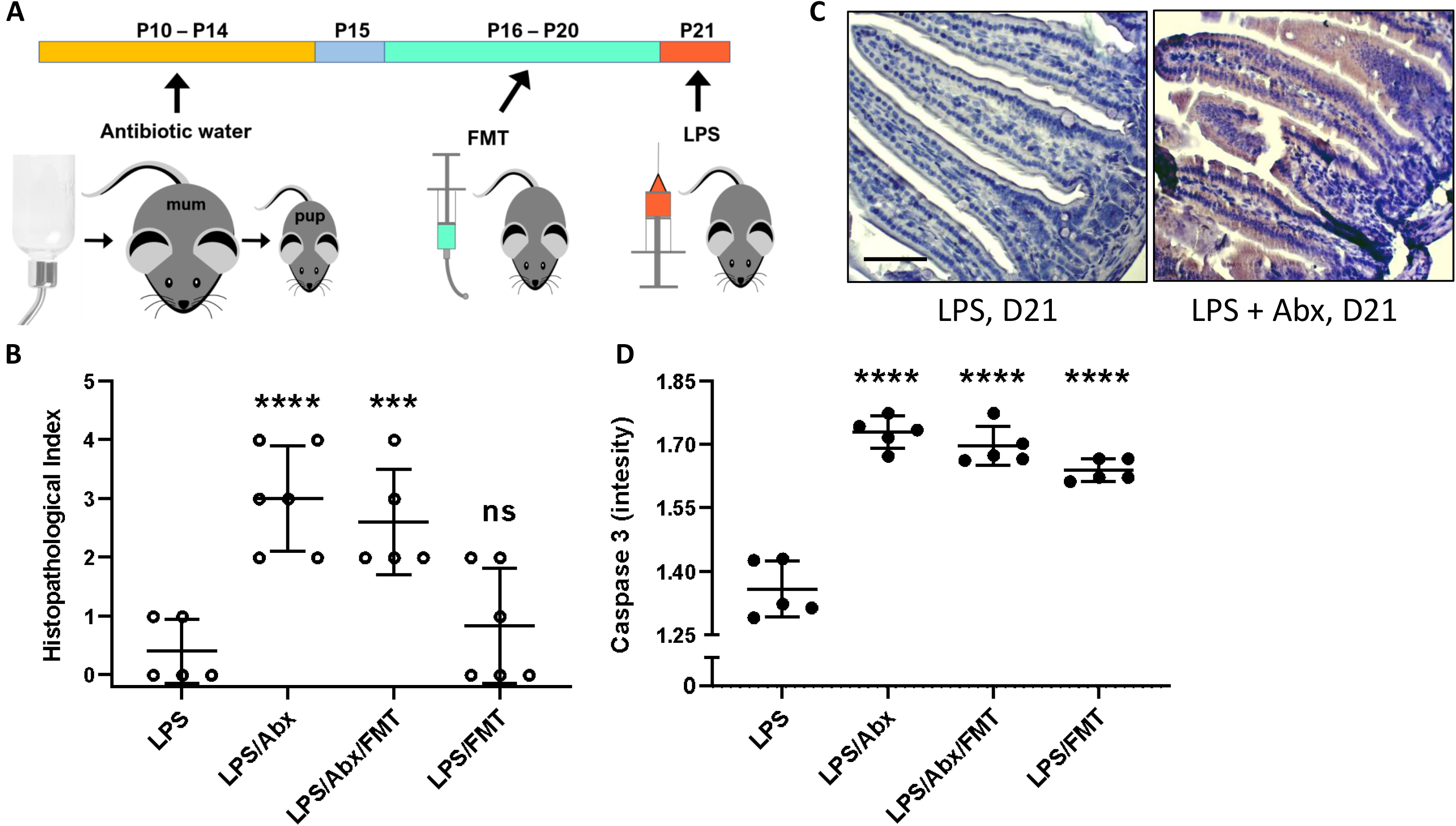
Neonatal mice have LPS shedding responses after FMT and antibiotic treatments. (**A**) FMT and antibiotic treatments were performed to disrupt the microbiota and assess the impact of LPS on cell shedding in D21 old mice (previously shown to be refractory to LPS-induced pathology). (**B**) Semi quantitative analysis of ileum by histopathology confirmed D21 mice were protected from damage caused by LPS. Neonatal mice that received FMT at D16-D20 showed no increase (P = 0.7587) in damage. Both Abx and Abx + FMT groups had significantly increased damage (p<0.0001 and 0.0007 respectively). (**C-D**) Apoptotic cell death was analysed using anti-cleaved caspase 3 (CC3). D21 mice had little to no apoptotic cell death which was significantly increased (P<0.0001) across all FMT and Abx dosed groups. Statistical analysis performed using one-way-Anova with Dunnett’s post-test n = 5. Scale bar 100 µm.

## Discussion

Using a mouse model of pathological intestinal cell shedding, we report that neonatal mice are largely refractory to LPS-induced epithelial cell death. Our data suggest a close association between the intestinal microbiota and developing immune system during the postnatal period that induces protection in this key developmental window. Moreover, disturbances of these pioneer microbial communities with antibiotics antagonises protective responses, highlighting the important role of the neonatal microbiota in regulating host physiology in health and disease.

The intestinal epithelium is the first line of defence between the luminal gut contents and the host. IEC integrity is maintained by a delicate balance of proliferation and cell shedding in the intestinal crypts and tips of the villi, respectively [8]. As such, regulation and maintenance of barrier integrity at the intestinal epithelial interface is vital to prevent entry of luminal microbes into the systemic circulation that may induce detrimental inflammatory responses. Indeed, there is a strong correlation between altered cell shedding and initiation of inflammatory conditions demonstrating the need to better understand the mechanistic links between cell shedding and disease progression [12]. Many intestinal inflammatory disorders present clinically in adolescents and adults, and we have previously shown that it is possible to dampen down pathological cell shedding in adult mice by supplementing with an early life microbiota member (i.e. *Bifidobacterium breve*) [16]. However, it has been suggested that early life events, which directly impact the infant/neonatal microbiota (such as diet and antibiotics), may shape the initiation of inflammatory cascades and IEC barrier integrity [22]. As such, addressing the role of the microbiota in regulating cell shedding in neonates is important and necessary to help develop our understanding of how these diseases evolve in children and adolescents.

To our surprise day-14-old mice were completely refractory to LPS-induced cell shedding, with limited cell shedding events in mice aged 21 days, and emergence of substantive cell shedding responses from day 29 postpartum. These data indicate that there is a ‘tipping’ point for induction of inflammatory cell shedding responses, which may correlate with the period when the microbiota becomes “adult-like”. Initially, we reasoned this reduced cell shedding may be linked to the operation of cellular signalling pathways that have previously been shown to modulate cell shedding response. However, neonates were competent in these aspects, including TNF-ɑ production, the functional cytokine that initiates apoptotic signalling cascades [19]. Nevertheless, we did note differential expression of other immune components in neonates (compared to adults), particularly IL-10 and IFN-Ɣ. IL-10 is a prototypical anti-inflammatory cytokine which inhibits the synthesis of pro-inflammatory cytokines such as TNF-ɑ and IFN-Ɣ thus limiting pathological cell shedding responses [23]. IL-10 induction has previously been described to be driven by certain microbiota members such as *Bacteroides*, *Bifidobacterium* and *Helicobacter*, suggesting that specific members may play a key role for inducing anti-inflammatory and anti-apoptotic responses, thereby driving protective barrier integrity during the neonatal period [24–27].

Our previous studies demonstrated an important role for infant-associated *Bifidobacterium breve*, which reduced pathological cell shedding processes in adult mice via the bifidobacterial exopolysaccharide capsule and host MyD88 signalling interactions [16, 28]. Although *Bifidobacterium* is found at high levels in infants, species and strains of this genus are not consistently detected in neonatal mice. Indeed, our microbiota profiling indicated very low levels of this taxa in our mice (0.7%), highlighting other early life bacterial members may be involved in the observed protective cell shedding responses at days 14 and 21. We observed lower microbial diversity in day 14 mice, when compared to their older counterparts, with correspondingly low concentrations of duodenum and stomach metabolites; similar to the relatively ‘simple’ infant microbial ecosystems [1]. Notably, we observed high relative abundance of typical early life facultative anaerobic species such as *Escherichia* and *Streptococcus* [3, 29]. Previous studies in germ-free and conventional mice have indicated *E. coli* plays a crucial role in immune programming, including signalling via *E. coli*-associated flagella (via TLR5). The intestinal epithelium represents a central interface for these two-way interactions with recent work showing that IEC expression of TLR5 is a critical step in promoting luminal colonisation of flagellated bacteria (e.g. *E. coli*); a window which closes at day 21 postpartum when TLR5 is down-regulated [30]. Other dominant neonatal microbiota members include Gram positive *Enterococcus* which has previously been shown (newborn-associated *Enterococcus faecalis*) to possess the ability to regulate certain cell signalling pathways, which results in transcriptional activation of IL-10 (increased in day 14 old mice) [31]. Crucially, as well as *Escherichia*, we also observed increased abundances of the Gram-negative Parabacteroides in neonatal mice. LPS also plays an important role in homeostatic immune modulation, alongside induction of pathological inflammatory responses as seen in our LPS-induced cell shedding model. Previous studies have indicated that immune cells are programmed to secrete inflammatory cytokines in response to LPS stimulation, and then become refractory to further stimulation (i.e. ‘exhausted’) [32]. Moreover, the order Bacteroidales (which includes *Parabacteroides*) produce a specific type of LPS (derived from an underacylated structural feature) that has been shown to silence TLR4 signalling, even when exposed to other gut microbiota member LPS types [33]. This suggests that the high relative abundance of these LPS-expressing bacteria may serve to downregulate pathological cell shedding responses and provide protection during the neonatal window. However, further studies would be required to test these hypotheses.

The day 21 postnatal period appears to show the most dynamic and variable microbial profile in mice, which may correlate with the ‘mixed’ nutritional exposure i.e. milk and mouse chow/coprophagy. In these day 21 mice, cell signalling responses induced at earlier time-points (by early life microbiota members) may help maintain epithelial barrier integrity, coupled with absence or very low levels of other microbiota members present in older mice, which may serve to override protective neonatal programming to allow typical pathological cell shedding. Weaning itself (and the start of coprophagia in mice) is associated with the emergence of a more adult-like microbiota. Notably, by day 29 postpartum, the intestinal microbiota showed greater similarity to the maternal community than at day 14 and day 21, with elevated relative abundance of *Candidatus* Arthromitus, *Alistipes* and *Lachnoclostridium*. Segmented Filamentous Bacteria (i.e. *Candidatus* Arthromitus) are often considered mouse-associated commensals, which can penetrate the intestinal mucus layer and intimately associate with IECs initiating immune responses, particularly inflammatory IL-17-associated responses [34, 35]. Moreover, facultative *Alistipes* spp. can overgrow in genetically susceptible mouse models which is correlated with induction of inflammatory colitis, thus linking presence of this taxa to the enhanced cell shedding responses observed in day 29 and adult mice [36]. Metabolomic profiling of the intestinal and stomach contents demonstrated emergence of a greater number of bacterial metabolites which coincided with the weaning period. These included microbial metabolites such as the SCFAs butyrate, propionate, and acetate. SCFAs can act either as substrates for host cell metabolism and/or as signalling molecules, particularly induction of immune signalling cascades via G-protein-coupled receptors (GPCRs) [37]. Previous studies have implicated SCFA-receptors signalling in chronic inflammatory diseases, which may correlate with the increased inflammatory responses observed in adult mice and enhanced pathological cell shedding [38]. Further SCFA supplementation studies are required to probe these findings.

Due to the significant differences in microbiota and metabolite profiles in the postnatal age groups, we hypothesised that disrupting the neonatal microbiota with clinically-relevant antibiotics would induce pathological cell shedding, thereby linking with studies that have shown that early life antibiotic usage correlates with increased risk of developing intestinal disorders [39]. Indeed, antibiotic exposure of neonatal mice prior to day 21 postpartum led to a significant increase in the cell shedding response. We also proposed that emergence of the adult like microbiota may represent the first, permissive step toward enabling cell shedding in this model. However, when neonatal mice were colonised with an adult microbiota (alone, without prior antibiotic treatment), we only observed enhancement of apoptosis (caspase 3), but not a concurrent increase in intestinal pathological further suggesting that it is the neonatal microbiota maintaining the epithelial barrier in response to inflammatory stimulus. Further work should seek to explore this early life window in greater detail to further interrogate the nature of the protective response, and the key microbial drivers and metabolites involved in this modulation [4].

In conclusion, our work has provided important new insights into the role of the early life intestinal microbiota in regulating the intestinal epithelium, barrier function and the immune system. It suggests a complex interplay between these compartments in protecting the neonatal gut from pathological cell shedding during the critical period of early life. These data may provide a platform for the development of therapies aimed at reducing chronic intestinal diseases in paediatric and adult patient populations.

## Materials and Methods

### Animals

All mouse experiments were performed under the UK Regulation of Animals (Scientific Procedures) Act of 1986. The project licences PDADA1B0C and PPL80/2545 under which these studies were carried out was approved by the UK Home Office and the UEA Animal Welfare Ethical Review Body. Time-mated female C57BL/6 Jax mice and adult C57BL/6 Jax mice (aged 6-10 weeks) were obtained from Charles River UK. All animals were maintained in the UEA Disease Modelling Unit under a standard 12 h light/dark cycle and fed standard rodent chow and water *ad libitum* under specific pathogen free (SPF) conditions.

### Induction of cell shedding and tissue collections

At 14, 21 and 29 days post-partum, pups (n = 6 per group) or adult control mice (n = 6) received an intra-peritoneal (IP) injection of 1.25mg/kg LPS from *Escherichia coli* 0111:B4 (Sigma), 0.5mg/kg murine Tumour Necrosis Factor Alpha (TNF-ɑ: 315-01A, Peprotech) or sterile saline (control). Mice were sacrificed 90 minutes post-challenge with LPS / TNF-ɑ. At autopsy, proximal small intestine was collected in 10% neutral buffered formalin (Sigma) and fixed for 24 h. Samples of proximal small intestine and spleen were also collected into RNA later for transcriptome analysis or frozen on dry ice for subsequent analysis (Figure 1A).

### Faecal microbiota transplant (FMT)

To determine the effects of the microbiota in the protection against LPS shedding we altered the gut microbiota with a combination of antibiotics and FMT from adult mice as per Table 1 and Figure 5A. At 10 days postpartum (D10) group 2 and 3 had 100 µg/ml ciprofloxacin and 500 µg/mL ampicillin added to the cage drinking water for 5 days to deplete the microbiota. After 5 days, both groups received normal drinking water for the rest of the study. For the FMT, two-three faecal pellets were collected from the control group Dam. Contents were gently homogenised using sterile glass beads and sterile PBS and passed through a 0.7 µm filter, with 50 µL of solution given to each pup by oral gavage. FMT’s were prepared daily, and pups were orally gavaged once a day from D16 to D20.

**Table 1:**
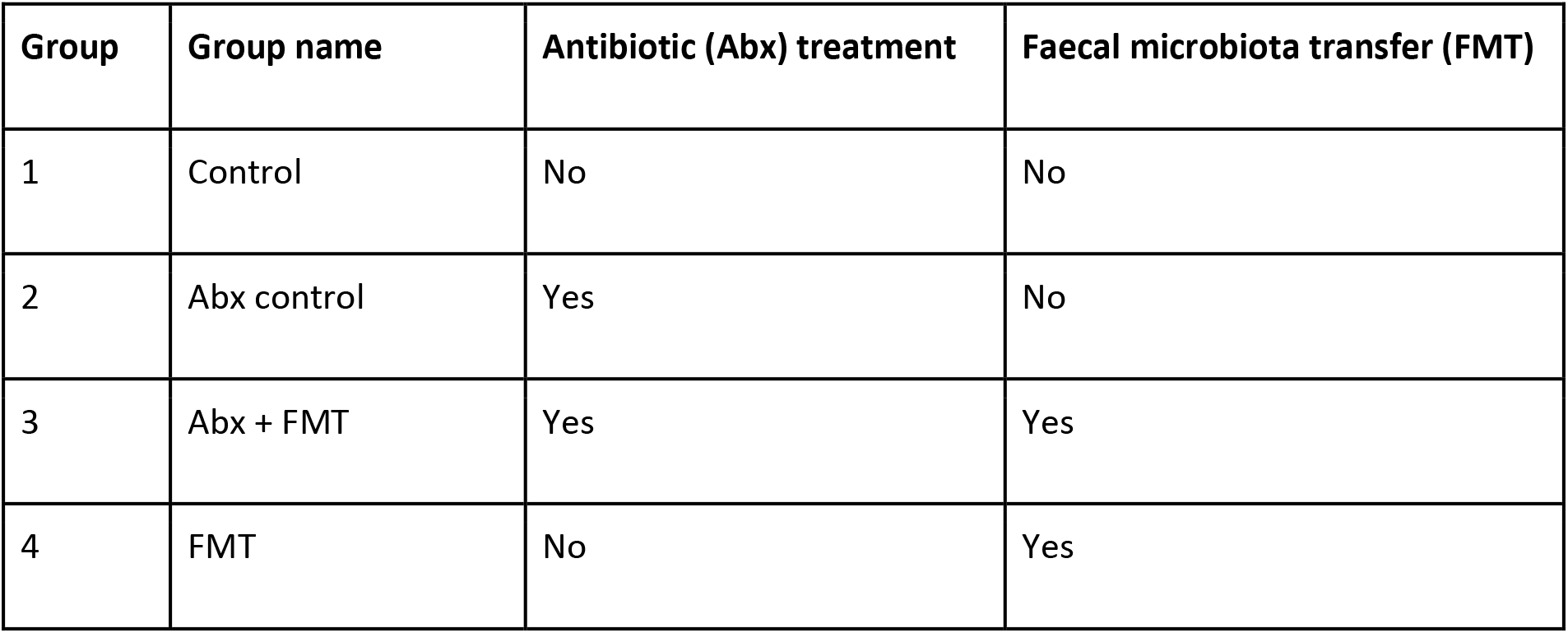
Treatment groups for FMT and antibiotic studies

### Histology

Formalin fixed tissue was dehydrated through a graded ethanol and xylene series and embedded in paraffin wax. 5 μm sections were used for immunohistochemistry. Following deparaffinization and rehydration, tissue sections were treated with 1% hydrogen-peroxide in methanol to block endogenous peroxidases. Subsequently, slides were treated using heat-induced antigen retrieval in 0.01M citrate acid buffer (pH 6) followed by antibody staining. CC3 staining was performed using rabbit polyclonal anti-active Caspase-3 antibody (AF835, R&D systems) and peroxidase-labelled anti-rabbit EnVision™ secondary antibody (Dako) or DAB or mouse polyclonal Caspase-3 antibody (NB100-56708H Novus Biologicals LTD) and visualised with DAB peroxidase (HRP) substrate kit (SK-4100, Vector Laboratories LTD). Rabbit polyclonal anti-active Caspase-3 antibody BrdU staining was performed using Sheep anti-BrdU-biotin (ab2284, Abcam) and neutravidin-HRP (31030, Pierce). TNF-R1 staining was performed using rabbit Anti-TNF-R1 antibody (sc-7895, Santa Cruz) and peroxidase-labelled anti-rabbit EnVision™ secondary antibody (Dako). Following secondary antibody binding, signal was detected using 3,3’-diaminobenzidine (Vector) followed by counterstaining with haematoxylin.

### Caspase-3 quantification

IECs were counted on a cell positional basis from villus tip (Cell position (CP) 1) down towards the crypts under 400x magnification. 20 well orientated hemi-villi were counted per mouse and analysed using the Score, WinCrypts (44) and PRISM (version 8) analysis software. IECs were defined as “normal” in cases where staining for active Caspase-3 was absent. Immuno-labelled cells with either unaltered or shedding morphology were treated as Caspase-3 positive. IEC’s stained with mouse polyclonal anti-active Caspase-3 antibody (NB100-56708H) were visualised with DAB and staining intensity was measured using FIJI (ImageJ 1.52h, National Institutes of Health, USA). Imaging was with an Olympus BX60 microscope and C10plus digital camera.

### ELISAs

Murine TNF-ɑ or Murine IL-10 Ready set go ELISA kits were used (eBioscience). Protocol was performed as per manufacturer’s instructions. In brief, Costar™ 9018 ELISA plates were coated with 100 µL/well of capture antibody diluted in Coating Buffer overnight at 4°C followed by blocking with ELISASPOT diluent. Small intestinal samples were homogenised in Cell Lytic MT buffer (Sigma) using Lysing matrix E beads (MP Biomedicals) and homogenates cleared by centrifugation at 10,000 rpm for 15 minutes. After quantification using BCA kit (Pierce), equivalent amounts of test samples or standard material were applied to wells in duplicate and incubated overnight at 4°C. Detection Antibody, Streptavidin-HRP and substrate were then applied and plates were read at 450 nm with 570 nm correction.

### Western blots

Spleens were homogenised in Cell Lytic buffer using Lysing matrix E beads (MP Biomedicals) and homogenates cleared by centrifugation at 10,000 rpm for 15 minutes. Homogenates were quantified by BCA analysis and 10 mg of total protein separated by SDS-PAGE with 10% acrylamide resolving gel. Proteins were then transferred to Hybond-P PVDF membrane (GE Healthcare). Immuno-staining was performed with 1/100 anti-MYD88 antibody (ab2064, Abcam) and 1/2000 Goat anti-Rabbit IgG HRP conjugate (Millipore) or 1/5000 Rabbit Anti-Beta-Actin antibody (ab8227, Abcam) and 1/5000 Goat anti-rabbit HRP. Signal was detected using Immobilon^TM^ Western chemiluminescent HRP substrate (Millipore) and the FluorChem E imaging system (Protein Simple).

### Cytokine antibody array

Small intestinal samples were homogenised in Cell Lytic MT buffer (Sigma) using Lysing Matrix E beads (MP Biomedicals) at a concentration of 50mg/ml. Homogenates were then cleared by centrifugation at 10,000 rpm for 15 minutes before quantification of protein using BCA kit (Pierce). 250 mg of total protein was then used to probe Murine antibody inflammation arrays (Abcam; ab133999) as per manufacturer’s instructions. In brief, membranes were for 30 min before application of homogenised whole small intestinal lysates in blocking buffer overnight at 4°C. Following washing, biotin conjugated anti cytokine antibodies were applied for 2 h before further washing and application of HRP conjugated streptavidin. Chemiluminescent detection was then performed using supplied detection reagents and visualization performed using FluorChem E imaging system (Protein Simple). See **Supplementary Table 1** for plate layout.

### Isolation of Intestinal epithelial cells

IECs were isolated using a modification of the Weisser method [40]. Briefly, whole small intestine was opened longitudinally, chopped into c.a. 5 mm^2^ pieces and washed in 0.154 M sodium chloride (NaCl) containing 1mM Dithiothreitol (DTT). Intestine was then incubated in a salt solution containing 1.5 mM Potassium Chloride (KCl), 96 mM NaCl, 27 mM Tri-sodium citrate (Na_3_C_6_H_5_O_7_), 8mM sodium di-hydrogen phosphate (NaH_2_PO_4_) and 5.6mM di-sodium hydrogen phosphate (Na_2_HPO_4_), pH 7.3 for 15 minutes at 37°C with gentle shaking (140rpm). Epithelial cells were then stripped from mucosa by further incubation in PBS containing 1.5mM Ethylene diamine tetraacetic acid (EDTA) and 0.5mM DTT for 15 minutes at 37°C, 140 rpm. Epithelial cells were then recovered and stabilised by pelleting at 1000 rpm for 5 minutes followed by lysis in buffer RLT RNA lysis solution (Qiagen).

### Isolation of DNA and 16S rRNA sequencing and analysis

Mouse faecal samples were collected into Lysis Matrix D tubes from the same mice at day 14, 21 and 29. Adult faecal samples were collected from both parents at the same time as day 14 samples. DNA extraction was carried out using a FastDNA Spin Kit for Soil (SKU 116560200-CF, MP Bioscience) following the manufacturers protocol with the exception of extending the bead-beating step to 3 minutes. DNA was quantified using a Qubit® 2.0 fluorometer (Invitrogen). 16S rRNA region (V1-V2) primers were used for library construction (**Supplementary Table 2**), with the following PCR conditions; cycle of 94°C 3 min and 25 cycles of 94°C for 45 s, 55°C for 15 s and 72°C for 30 s. Sequencing of the 16S rRNA gene libraries was performed using Illumina MiSeq platform with 300 bp paired end reads. Raw reads were filtered through quality control using trim galore (version 0.4.3), minimum quality threshold of phred 33, and minimum read length of 60 bp. Reads that passed threshold were aligned against SILVA database (version: SILVA_132_SSURef_tax_silva) using BLASTN (ncbi-blast-2.2.25+; Max e-value 10e-3) separately for both pairs. After performing BLASTN alignment, all output files were imported and annotated using the paired-end protocol of MEGAN on default Lowest Common Ancestor (LCA) parameters [41]. For analysis of microbiota composition, the samples were normalised by subsampling each sample to 3224 reads (the number of reads of the lowest sample) to equalize library sizes using the Phyloseq package version 1.24.2 in RStudio version 1.1.463.

### Isolation of RNA, cDNA synthesis and real-time PCR

RNeasy plus mini spin columns were used to isolate total RNA (Qiagen). In brief, samples were homogenised using a rotor stator hand-held homogeniser in buffer RLT (tissue samples) or reconstituted in buffer RLT by pipetting (isolated IECs). Lysates were processed through QIAshreddor columns (Qiagen) and subsequently RNeasy mini-spin columns. Purified RNA was eluted into RNAase free water.

For TNF-R1 real-time PCR analysis of isolated IECs, cDNA was synthesized using Quantitect reverse transcription kit (Qiagen) and real-time PCR performed using Quantitect SYBR green mastermix (Qiagen) and Quantitect murine TNF-R1 primer assay (Qiagen) or hypoxanthine– guanine phosphoribosyltransferase (HPRT) 5′-GACCAGTCAACAGGGGACAT-3′ (sense) and 5′-AGGTTTCTACCAGTTCCAGC-3′ (antisense) primers. Cycling was performed on a Roche LightCycler 480 using the following conditions: 95°C, 5 min then 40 cycles of 95°C, 10 s; 60°C, 35 s. Relative quantification of levels of transcript expression was calculated using the Pfaffl method by comparing cycle threshold (CT) value of each target gene to the CT value of housekeeper [42]. Data are presented as a ‘fold change’ in expression (normalized against control untreated mice per cells).

For PCR array analysis, reverse transcription was performed using the Transcriptor first strand cDNA synthesis kit (Roche) and cDNA analysed on custom real-time ready 384 well custom PCR arrays (Roche). Arrays contained the following murine targets: IL-1B, Ml2, Arrdc2, Bid, Card6, Casp4, Cycs, Ikbe, Rel, Tlr2, Traf5, Tnfrs8, Tnfrsf1b, Tnfrs19, Bd2l1, Rn18s, Il6, Alpi, Pdzd7, Casp12, Card11, Casp7, Fadd, Nkb1, Rela, Traf6, Traf2, Tnfrs9, Tnfrs11b, Tnfrs15, Bd2l10, Rpl13a, Il10, Pdx1, Apaf1, Card10, Casp3, Casp9, Fas, Nkb2, Relb, Traf3, Traf7, Tnfsf11, Tnfsf25, Tnfsf21, Bfar, Ptgds, Ifng, Ceacam10, Bad, Casp1, Caps6, Cd40lg, Fasl, Nkbia, Serpinb2, Tnf, Tnfsf18, Tnfsf8, Tnfsf4, Tnfsf14, Ephb3, Ptgs2, Col18a1, Pzp, Bax, Card9, Casp2, Cd40, Faslk, Nkbib, Serpinb9, Traf1, Tnfsf10, Tnfrsf10b, Tnfrsf1a, Tnfsf18, Hprt1, Ddk1, Gasdma, Fgl1, Bd2, Casp14, Casp8, Cflar, Ikbkb, Nod2, Actb, B2m, Gapdh Cycling was performed using Lightcycler 480 probe master mix and a Roche LightCycler 480 using the following conditions: 95°C, 10 min then 40 cycles of 95°C, 10 s; 60°C, 30 s, 72°C, 1 s. Final analysis of CT values was using the DDCT method as described above.

### NMR metabolomics

Stomach contents were collected from mice at autopsy and immediately frozen on dry ice before transfer to −80°C prior to analysis. Small intestinal contents were flushed from the duodenum using sterile water and frozen similarly prior to analysis.^1^H NMR was used to identify the presence and concentration of several metabolites. Supernatant samples were thawed at room temperature and prepared for ^1^H NMR spectroscopy by mixing ~50 mg (FW) of stomach content samples with 600*µ*L NMR buffer (0.26 g NaH_2_PO_4_ and 1.41 g K_2_HPO_4_) made up in 100% D_2_O (100 ml), containing 0.05% NaN_3_ (50 mg), and 1 mM sodium 3-(Trimethylsilyl)-propionate-*d*4, (TSP) (17 mg) as a chemical shift reference. The sample was mixed, and 500 *µ*L was transferred into a 5-mm NMR tube for spectral acquisition. The ^1^H NMR spectra were recorded at 600MHz on aBruker Avance spectrometer (Bruker BioSpin GmbH, Rheinstetten, Germany) running Topspin 2.0 software and fitted with a 5 mm TCI cryoprobe. Sample temperature was controlled at 300 K. Each spectrum consisted of 512 scans of 65,536 complex data points with a spectral width of 12.3 ppm (acquisition time 2.66 s). The noesypr1d pre-saturation sequence was used to suppress the residual water signal with low power selective irradiation at the water frequency during the recycle delay (D1 = 3 s) and mixing time (D8 = 0.01 s). A 90° pulse length of 12.9 μs was set for all samples. The “noesypr1d” pre-saturation sequence was used to suppress the residual water signal with a low-power selective irradiation at the water frequency during the recycle delay. Spectra were transformed with a 0.3-Hz line broadening, manually phased, baseline corrected, and referenced by setting the TSP methyl signal to 0 ppm. The metabolites were quantified using the software Chenomx NMR suite 7.6™. Metabolites were identified using information found in the literature (references) or on the web (Human Metabolome Database, http://www.hmdb.ca/) and by use of the 2D-NMR methods, COSY, HSQC, and HMBC.

### Statistics

Results were plotted and analysed using PRISM version 8 software (GraphPad). Data (mean + s.d.) were analysed using ordinary one-way anova with Tukeys post-test with a 95% confidence interval. CC3 was measured using FIJI (Fiji Contributions) and plotted as mean fluorescence intensity (MFI) from 3 sections per mouse (n = 7). Microbiota and metabolome data analysis was carried out in R version 3.5.0 using RStudio (version 1.1.463). The ‘vegdist’ function in the ‘vegan’ (version 2.5.4) R package was used to calculate Bray-Curtis dissimilarity matrices for the microbiota abundance and metabolomics data. NMDS plots of Bray-Curtis matrices were generated for microbiota data using the vegan package. Permutational multivariate analysis of variance (PERMANOVA) was performed on Bray–Curtis matrices using the ‘pairwiseAdonis’ package. Diversity analysis of microbial abundance data was performed using the ‘vegan’ package. Statistical analysis of number of genera and Shannon diversity between groups was performed using Kruskal–Wallis one-way analysis of variance with Pairwise Wilcoxon Rank Sum Tests using Benjamini–Hochberg stepwise adjustment.

## Author contributions

KRH and LJH designed the study. KRH and ZS performed the animal studies, downstream sample processing, and associated experiments. LC processed the faecal samples and sequencing library. SC processed the 16S rRNA sequencing data. FB performed western blots. CA-G helped with histology and caspase-3 quantification. GL-G carried out the NMR experiments. KRH, ZS, MJD and LJH analysed and visualised the data. KRH, ZS and LJH drafted the manuscript with further interpretation of data and drafting from MJD and AJMW. All authors read, gave comments, and approved the final manuscript.

## Competing interests

The authors declare that they have no competing interests.

## Acknowledgments

This work was funded via a Wellcome Trust Investigator Award to LJH (100974/Z/13/Z) and support of the BBSRC Norwich Research Park Bioscience Doctoral Training Grant (BB/M011216/1, supervisor LJH, student CAG), Institute Strategic Programme (ISP) grant for Gut Health and Food Safety, BB/J004529/1 (AJMW and LJH), and ISP grant for Gut Microbes and Health BB/R012490/1 and its constituent project(s), BBS/E/F/000PR10353 and BBS/E/F/000PR10355 to LJH.

## Notes

#### Summary of Updates

Update to author name (Kevin R Hughes)

