## Supplementary Files for "The early life microbiota protects neonatal mice from pathological small intestinal epithelial cell shedding"

| | A | B | C | D | E | F | G | H | I | J | K | L |
|---|---|---|---|---|---|---|---|---|---|---|---|---|
| 1 | **Pos Cntrl** | **Pos Cntrl** | **Neg Cntrl** | **Neg Cntrl** | **BLANK** | **BLC** | **CD30 L** | **Eotaxin** | **Eotaxin-2** | **Fas Ligand** | **Fractalkine** | **GCSF** |
| 2 | **Pos Cntrl** | **Pos Cntrl** | **Neg Cntrl** | **Neg Cntrl** | **BLANK** | **BLC** | **CD30 L** | **Eotaxin** | **Eotaxin-2** | **Fas Ligand** | **Fractalkine** | **GCSF** |
| 3 | **GM-CSF** | **GM-CSF** | **IL-1α** | **IL-1β** | **IL-2** | **IL-7** | **IL-4** | **IL-6** | **IL-9** | **IL-10** | **IL-12 p40/p70** | **IL-12 p70** |
| 4 | **GM-CSF** | **GM-CSF** | **IL-1α** | **IL-1β** | **IL-2** | **IL-7** | **IL-4** | **IL-6** | **IL-9** | **IL-10** | **IL-12 p40/p70** | **IL-12 p70** |
| 5 | **IL-13** | **IFN-γ** | **I-TAC** | **KC** | **Leptin** | **LIX** | **Lymphotactin** | **MCP-1** | **MCSF** | **MIG** | **MIP-1α** | **MIP-1γ** |
| 6 | **IL-13** | **IFN-γ** | **I-TAC** | **KC** | **Leptin** | **LIX** | **Lymphotactin** | **MCP-1** | **MCSF** | **MIG** | **MIP-1α** | **MIP-1γ** |
| 7 | **RANTES** | **SDF-1** | **TCA-3** | **TECK** | **TIMP-1** | **TIMP-2** | **TNF-α** | **sTNF RI** | **sTNF RII** | **BLANK** | **BLANK** | **Pos Cntrl** |
| 8 | **RANTES** | **SFD-1** | **TCA-3** | **TECK** | **TIMP-1** | **TIMP-2** | **TNF-α** | **sTNF RI** | **sTNF RII** | **BLANK** | **BLANK** | **Pos Cntrl** |

**Supplementary Table 1**: Antibody array layout

A

B

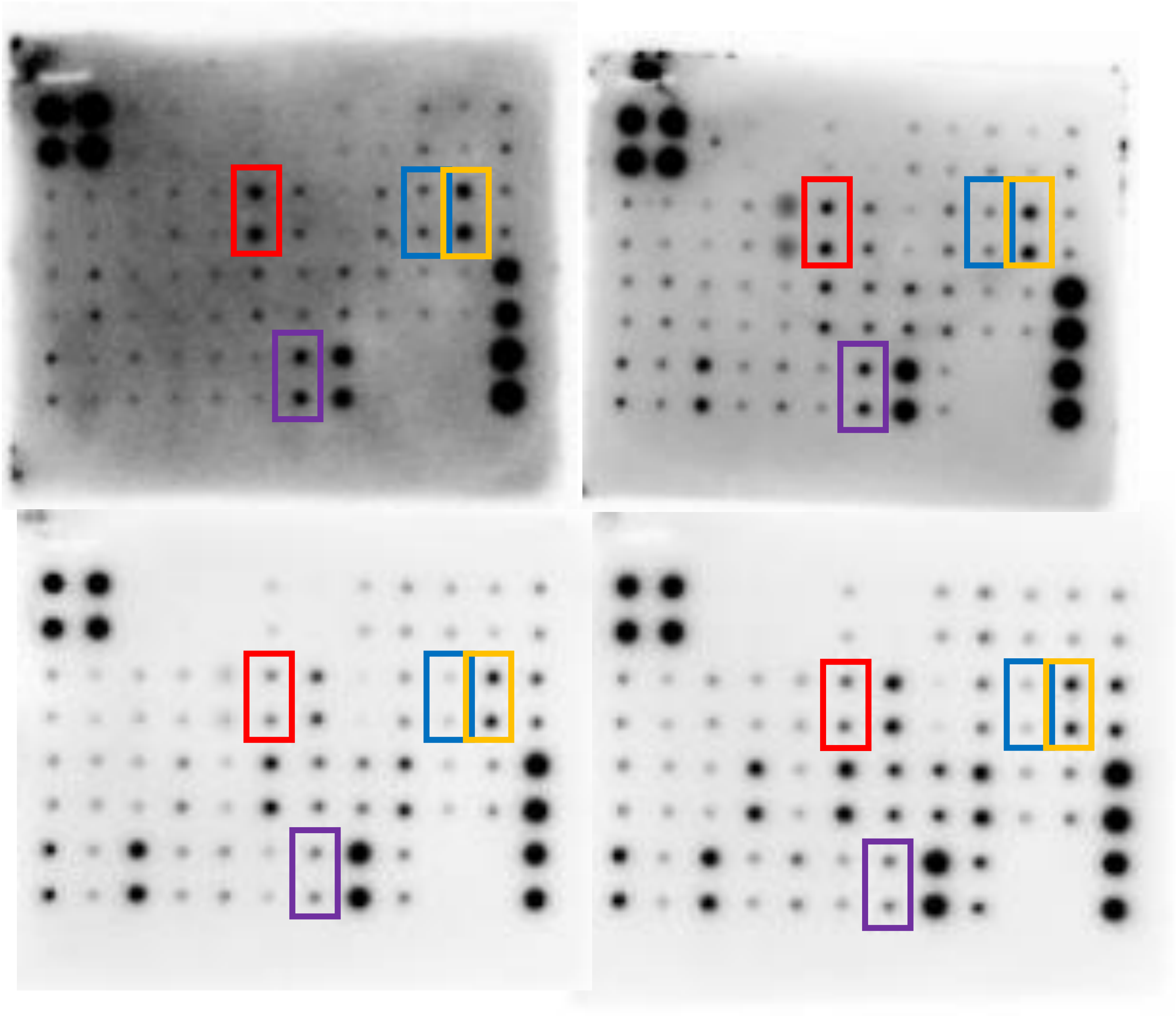

**Supplementary Figure 1: Neonatal mice have altered cytokine responses**
Antibody arrays were used to assess cytokine changes in neonatal versus adult homogenised small intestinal samples (see **Supplementary Table 1** for full play layout). (**A**) Two experimental replicate antibody arrays from D14 neonates after treatment with 1.25mg/kg LPS. (**B**) Adult (8-10 week old) antibody arrays after LPS treatment. Coloured boxes highlight elevated differences in neonatal homogenates vs. adult samples; red (IL-7), blue (IL-10), yellow (IL-12), and purple (TNF-α).

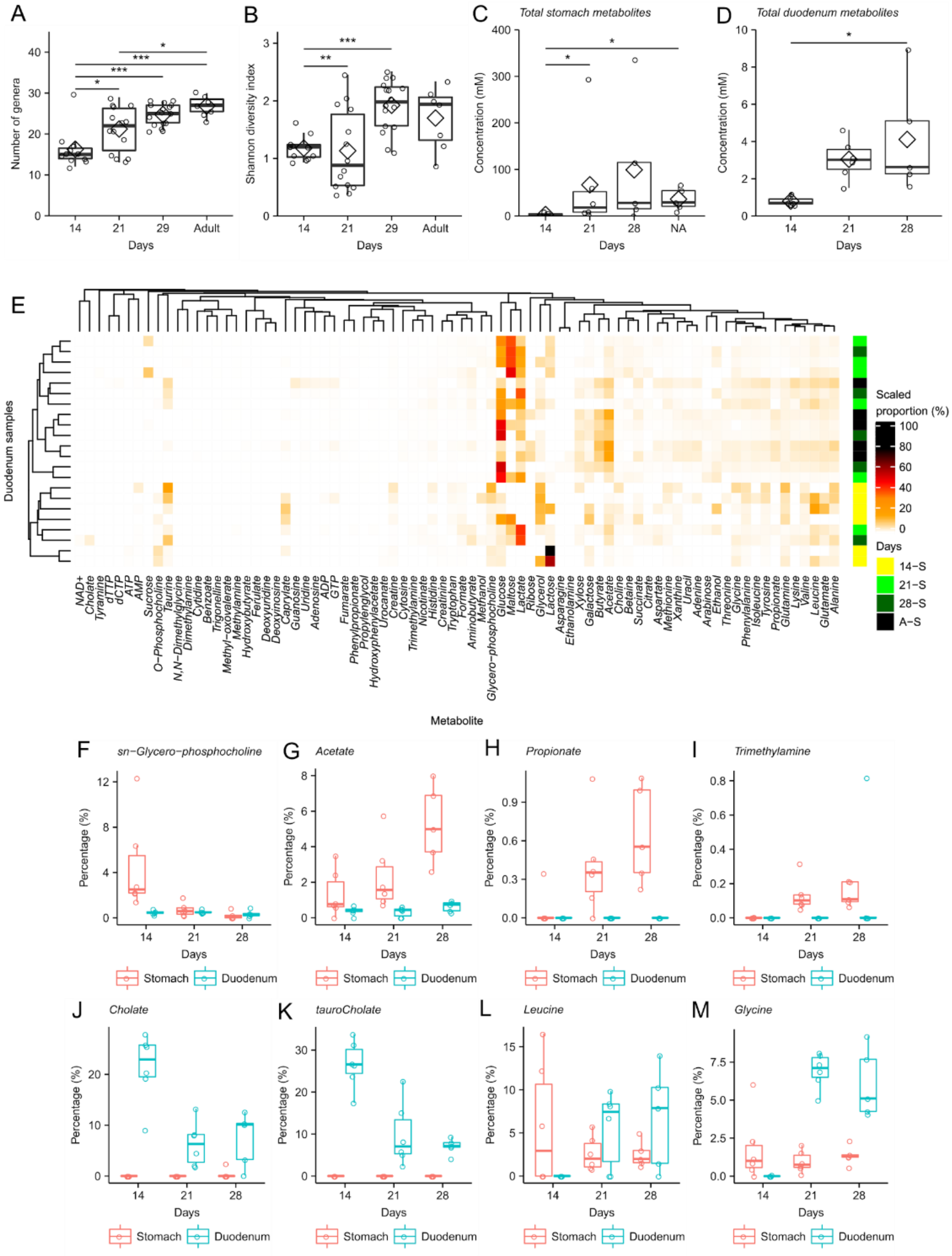


**Supplementary Figure 2: Microbiota diversity, total metabolites and stomach metabolite composition.** (**A**) Number of genera detected in each faecal sample. (**B**) The Shannon Diversity index of the microbiota in each sample at the genus level. (**C**) The total concentration of metabolites in stomach contents. (**D**) The total concentration of metabolites in duodenum contents. (**E**) Heatmap showing proportions of metabolites in stomach contents determined using NMR analysis and clustered using a Bray-Curtis matrix shows distinct grouping of metabolites at mice aged 14 days. (**F**) sn-Glycero-phosphocholine, (**G**) Acetate, (**H**) Propionate, (**I**) Trimethylamine, (**J**) Choline, (**K**) tauroCholine, (**L**) Leucine, and (**M**) Glycine. Boxplots show mean (diamond), median (solid bar), and interquartile range while circles show individual samples. Statistical significance determined by ANOVA with post hoc Tukey test or Kruskal–Wallis test with post hoc Pairwise Wilcoxon Tests. *= $p < 0.05$, **= $p < 0.01$, and ***= $p < 0.001$.
